# International risk of yellow fever spread from the ongoing outbreak in Brazil

**DOI:** 10.1101/152280

**Authors:** I Dorigatti, A Hamlet, R Aguas, L Cattarino, A Cori, CA Donnelly, T Garske, N Imai, NM Ferguson

## Abstract

The largest Yellow Fever (YF) outbreak in a decade in Latin America is underway in the Southeast of Brazil. In this article we provide a quantitative assessment of the risk of travel-related international spread of YF. We argue that mitigating the risk of imported YF cases seeding local transmission requires heightened surveillance in the southern United States, Latin America (especially Argentina, Chile and Uruguay) and Europe (especially Portugal, Spain, Italy and Germany).

## Introduction

The largest Latin American outbreak of Yellow Fever (YF) reported in a decade is currently unfolding in Brazil, with 784 confirmed human cases and 267 confirmed deaths reported as of 31^st^ May 2017 [1] (Figure A-B). The epidemic has spread from Minas Gerais and Espírito Santo to São Paulo and Rio de Janeiro, thus raising public health concern about the establishment of urban transmission and the spread of YF beyond Brazil’s national border.

By linking the latest epidemiological data [1] with World Tourism Organization data on the volume of air, land and water border crossings [2], we assessed the risk of travel-related international spread of YF.

### Data Sources

The cumulative number of confirmed cases reported in the Southeast of Brazil was obtained from the weekly epidemiological bulletins on YF published online by the Brazilian Ministry of Health [3]. The data used in this analysis refer to bulletin number 43 of 31^st^ May 2017 [1].

For each state, information on the first and last dates of symptom onset of confirmed cases (Table) has been retrieved from the time series reported in bulletin number 43 of 31^st^ May 2017 [1] using a web plot digitalizer tool [4].

Country- and state-level population data for Brazil relative to 2016 were obtained from the Brazilian Institute of Geography and Statistics website [5].

Data on the annual volumes of air, land and water border crossings for Brazil relative to inbound (arrivals of non-resident tourists at national borders by country of residence) and outbound (trips abroad by Brazilian resident visitors to countries of destination) tourism for year 2015 have been purchased from the World Tourism Organization (UNWTO) [2].

Information on the monthly distribution of inbound tourism and on the average duration of stay of international visitors to Brazil by country of origin has been obtained from a survey on the touristic demand in Brazil conducted in 2015 [6].

**Table:** Dates of first and last symptom onset of confirmed YF cases reported in the Southeast Brazil as of 31^st^ May 2017 [1].

| State | First date of symptom onset | Last date of symptom onset |
| --- | --- | --- |
| Minas Gerais | 19/12/2016 | 20/04/2017 |
| Espírito Santo | 04/01/2017 | 30/04/2017 |
| São Paulo | 17/12/2016 | 20/04/2017 |
| Rio de Janeiro | 19/02/2017 | 10/05/2017 |

## Methods

We estimated the expected number of YF cases departing Brazil before recovery, comprising infected Brazilian residents travelling abroad during the incubation or infectious periods (“exportations”) and international tourists infected by YF during their stay in the Southeast of Brazil and returning to the home country before the end of the infectious period (“importations”).

### Exportations

Let *C_s,w_* denote the cumulative number of confirmed YF cases reported in state in time window *W*, with *W* denoting the number of days between the first and the last confirmed YF case in state *S*

Comparison of the observed case fatality ratio (CFR) among confirmed cases in Brazil (34.1%) with the established CFR among severe cases (47%, 95% CI 31 – 62%), mild or severe cases (13%, 95% CI 5 – 28%) and YF infections (5%, 95% CI 2 – 12%) [7] suggests that case confirmation in the 2017 YF outbreak in Brazil is broadly consistent with severe cases being detected. Therefore we assumed that all confirmed cases are severe and, following Johansson et al. [7], that there are 9 mild or asymptomatic infections for each severe case. This implies that the cumulative number of YF cases in state *S* in time window*W* is given by 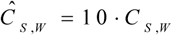

Let *pop_s_* denote the resident population of state *S*, *pop_B_* denote the resident population of the whole of Brazil and *T_D_* denote the annual number of Brazilian travellers visiting country *D*. The per capita probability that a Brazilian resident travels to country *D* during time window *W* is given by 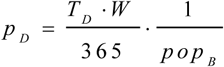.

We assumed that the incubation period *T_E_* is log-normally distributed with mean 4.6 days and variance 2.7 days [8] and that the infectious period *T_I_* is normally distributed with mean 4.5 days and variance 0.6 days [9]. The probability *P_i_* that a YF case incubates or is infectious in time window *W* is 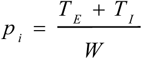.

The number of residents of state *S* infected by YF and travelling abroad during the incubation or infectious periods in time window *W* is given by 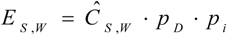.

Variability in the incubation and infectious periods were accounted for by sampling 10,000 times *T_E_* and *T_I_* from their respective distributions, leading to a full distribution for *P_i_* and in turn for *E_S,W_*.

### Importations

Let *T_O_* denote the annual number of travellers visiting Brazil from country *O*, *f_m_* the proportion of international travellers visiting Brazil in month *m* and *p_s,m_* the relative proportion of the epidemic window *W* In state *S* occurring in month *m*. Assuming that travellers to Brazil pick destination states within the country with a probability proportional to the states’ population sizes, the expected number of travellers 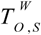 visiting state *S* from country *O* in in time window *W* is given by 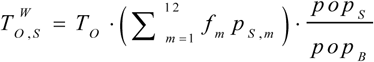.

Let *L_O_* denote the average length of stay of travellers visiting Brazil from country *O*. The per capita risk of infection of travellers visiting state *S* during their stay is estimated as 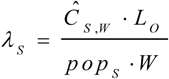.

The probability of returning to the home country while incubating or infectious is 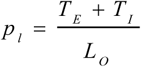, where *p_i_* is set to 1 if (*T_E_* + *T_I_*) > *L_O_*.

The expected number of international tourists infected by YF during their stay in state *S* and returning to the home country *O* before the end of the infectious period is estimated by 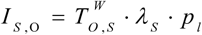.

Variability in the incubation and infectious periods were accounted for by sampling 10,000 times *T_E_* and *T_I_* from their respective distributions, leading to a full distribution for *p_l_* and in turn for *I_s,o_*.

## Results and Discussion

We show in Figure C the expected number of YF cases departing Brazil before recovery, comprising exportations and importations, for the countries with an upper 95% limit exceeding 1 case over all states. The United States and Argentina, which contain regions potentially suitable for YF transmission, may have already received at least one travel-related YF case capable of seeding local transmission. Europe (especially Portugal, Spain, Italy and Germany) and Latin America (especially Chile and Uruguay) are also at risk of YF importation. Increased awareness, monitoring and preparedness is therefore appropriate to avoid the current YF outbreak in Brazil seeding new YF outbreaks globally.

**Figure:**
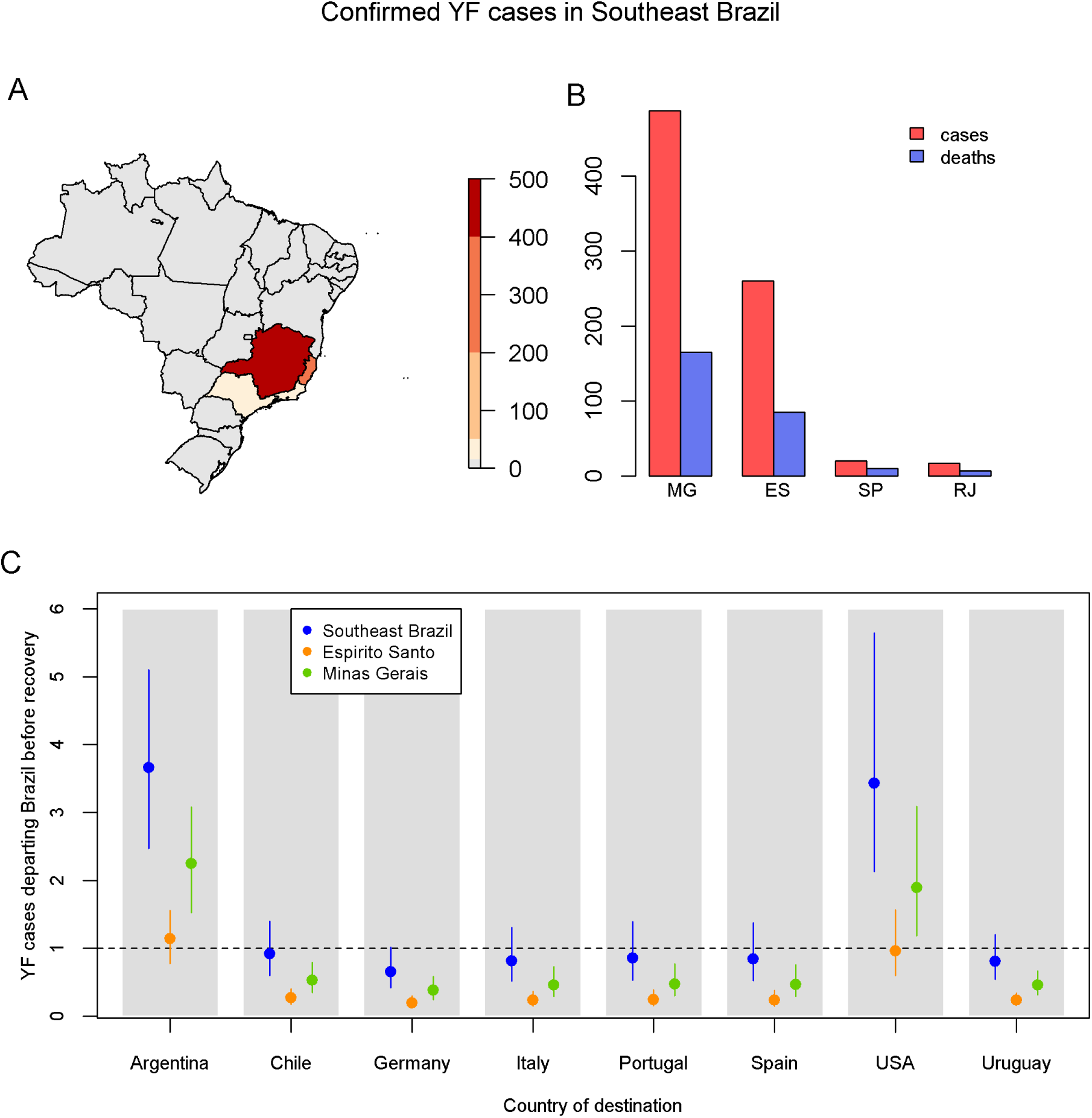
A) Geographical distribution of the total number of confirmed cases reported as of 31^st^ May 2017 in the Southeast of Brazil since December 2016 [1]; B) Total number of confirmed cases and confirmed deaths by state reported as of 31^st^ May 2017 [1]. MG stands for Minas Gerais, ES stands for Espírito Santo, SP stands for São Paulo and RJ stands for Rio de Janeiro; C) Mean and 95% confidence interval of the estimated number of YF cases that could potentially seed a YF outbreak in the destination countries, comprising infected Brazilian residents travelling abroad during the incubation or infectious period (exportations) and international tourists infected by YF during their stay in the Southeast of Brazil and returning to the home country before the end of the infectious period (importations). The mean and 95% confidence intervals have been obtained by numerically sampling 10,000 times the incubation and infectious period distributions [8,9]. Only countries with an upper 95% limit exceeding 1 exported case over all states (Southeast Brazil) are shown. The estimated risk of international spread from São Paulo and Rio de Janeiro is minimal and is not shown. USA stands for the United States of America.

## References

1. Monitoramento dos casos e óbitos de febre amarela no Brasil, informe n. 43/2017, accessed on 16^th^ June 2017 at http://portalarquivos.saude.gov.br/images/pdf/2017/junho/02/COES-FEBRE-AMARELA---INFORME-43---Atualiza----o-em-31maio2017.pdf

2. World Tourism Organization (2016), Yearbook of Tourism Statistics dataset [Electronic], UNWTO, Madrid, data updated on 27/04/2016.

3. http://portalsaude.saude.gov.br/index.php/o-ministerio/principal/leia-mais-oministerio/619-secretaria-svs/l1-svs/27300-febre-amarela-informacao-e-orientacao

4. B. Tummers, DataThief III. 2006 http://datathief.org/

5. Data accessed on 23^rd^ May 2017 at http://downloads.ibge.gov.br/downloadsestatisticas.htm

6. Estudo da Demanda Turística Internacional – 2015 accessed on 12^th^ May 2017 at http://www.dadosefatos.turismo.gov.br/2016-02-04-11-54-03/demanda-tur%C3%ADsticainternacional.html

7. Johansson MA, Vasconcelo PFC and Staples JE, The whole iceberg: estimating the incidence of yellow fever virus infection from the number of severe cases, Trans R Soc Trop Med Hyg. 2014; 108: 482–487

8. Johansson MA, Vasconcelo PFC and Staples JE, Incubation periods of yellow fever virus, Am J Trop Med Hyg. 2010; 83(1) 183–188

9. Monath TP, Yellow fever: an update. Lancet Infect Dis. 2001; 1(1):11-20

